# Interactive phenotyping of large-scale histology imaging data with HistomicsML

**DOI:** 10.1101/140236

**Authors:** Michael Nalisnik, Mohamed Amgad, Sanghoon Lee, Sameer H. Halani, Jose Velazquez Vega, Daniel J Brat, David A Gutman, Lee AD Cooper

**Affiliations:** Department of Biomedical Informatics, Emory University School of Medicine; Department of Neurology, Emory University School of Medicine; Emory University School of Medicine; Department of Pathology & Laboratory Medicine, Emory University School of Medicine; Winship Cancer Institute, Emory University; Department of Biomedical Engineering, Georgia Institute of Technology / Emory University School of Medicine Atlanta GA, USA

## Abstract

Whole-slide imaging of histologic sections captures tissue microenvironments and cytologic details in expansive high-resolution images. These images can be mined to extract quantitative features that describe histologic elements, yielding measurements for hundreds of millions of objects. A central challenge in utilizing this data is enabling investigators to train and evaluate classification rules for identifying objects related to processes like angiogenesis or immune response. Here we present HistomicsML, an interactive machine-learning framework for large whole-slide imaging data. HistomicsML uses active learning direct user feedback, making classifier training efficient and scalable in datasets containing 10^8^+ histologic objects. We demonstrate how HistomicsML can be used to phenotype microvascular structures in gliomas to predict survival, and to explore the molecular pathways associated with these phenotypes. Our approach enables researchers to unlock phenotypic information from digital pathology datasets to investigate prognostic image biomarkers and genotype-phenotype associations.

## INTRODUCTION

Slide scanning microscopes can digitize entire histologic sections at 200X-400X magnification, generating expansive high-resolution images containing 10^9^+ pixels. For cancer tissues, these images contain important biologic and prognostic information, capturing the diverse cytologic elements involved in angiogenesis, immune response, and tumor/stroma interactions. Image analysis algorithms can mine whole-slide images to delineate objects like cell nuclei, and to extract 10s-100s of quantitative features that describe the shape, color, and texture of each object. These histology-omic or “histomic” features can be used to train machine-learning algorithms to classify important elements like tumor-infiltrating lymphocytes, vascular endothelial cells, or fibroblasts. Identifying these elements in tissues requires considerable expertise, and imparting this knowledge to algorithms enables precise characterization of large imaging datasets in ways not possible by subjective visual assessment. Quantitative measures of the abundance, morphologies and spatial patterns of these elements can help investigators understand relationships between histologic phenotypes and survival, treatment response, and underlying molecular mechanisms. Studies that generate whole slide images can yield histomic annotations of 10^8^+ objects, and a central challenge in utilizing this data is in enabling domain experts to train classification rules and to evaluate their accuracy. With each image containing up to 10^6^+ discrete objects, facilitating interaction with domain experts requires fluid navigation of gigapixel images, visualization of derived image segmentation boundaries, mechanisms to intelligently acquire training data from experts, and to visualize classifications for millions of objects.

Histopathology image analysis has received significant attention with algorithms having been developed to predict metastasis (1), survival (2–6), grade (7, 8), and histologic classification (9–11), and to link histologic patterns with genetic alterations or molecular disease subtypes (12–14). Many algorithms demonstrate scientific or potential clinical utility, but few directly engage domain experts in analyzing histomic data (15, 16). Inputs are typically acquired offline by presenting a small collection of manually selected image sub regions to an expert for labeling or annotation. Enabling experts to directly interact with machine learning algorithms on large datasets creates a feedback loop that has been shown to improve prediction accuracy and user experience in general applications (17–21). In this feedback paradigm, the expert iteratively improves a classification rule by correcting or confirming predictions on unlabeled examples, cycling between labeling and training and prediction. Active learning extends this paradigm by identifying and labeling the examples that provide the most benefit to the classifier in each cycle. This approach seeks to increase the diversity of labeled examples used for classifier training, and avoids labeling redundant examples that are unlikely to improve performance. The challenge in utilizing active learning with histomic data is in building software with the scalable visualization and machine-learning capabilities described above.

This paper describes the histomics machine-learning system (HistomicsML), a software framework for interactive classification and phenotyping of large whole-slide imaging datasets. HistomicsML has several key features that enables users to rapidly train accurate histologic classification rules: (i) A web-based interface that fluidly displays gigapixel images containing 10^6^+ image analysis objects (ii) A machine-learning server for fast, interactive training of classification rules (iii) Active-learning algorithms for improved training efficiency and accuracy (iv) Tools for creating, sharing and reviewing labeled data and ground-truth validation sets. HistomicsML is open-source (https://github.com/cooperlab/ActiveLearning) and available as a software container for easy deployment (https://hub.docker.com/r/histomicsml/active/). Using imaging, clinical and genomic data from The Cancer Genome Atlas (TCGA), we demonstrate how HistomicsML can be used to develop an accurate classifier of vascular endothelial cell nuclei with minimal effort. We use this classifier to describe the phenotypes of microvascular structures in gliomas, and show that these phenotypes predict survival independent of both grade and molecular subtype. Finally, we identify molecular pathways associated with disease progression through integrated pathway analysis of mRNA expression data.

## RESULTS

### Active learning classification software for histology imaging datasets

An overview of the HistomicsML software is presented in Figure 1. Image segmentation algorithms are used to delineate histologic objects like cell nuclei in whole-slide images, and a histomic feature profile is extracted to describe the shape, texture, and staining characteristics of each delineated object (see Supplementary Figure 1). Images, features, and object boundaries are stored in a database and disk array to support visualization and machine-learning analysis. A web-browser interface enables users to rapidly train classification rules and review their predictions in large datasets containing 10^8^+ objects. A multiresolution image viewer provides zooming and panning of gigapixel images and dynamically displays object boundaries. A caching and pre-fetching strategy is used to display boundaries for objects in the current field of view and to fluidly handle panning events (see Supplementary Figure 2). Boundaries are color-coded to indicate their predicted class (e.g. green - endothelial cell nuclei) or membership in the training set (e.g. yellow – labeled). Users can refine the classification rule by clicking objects in the viewport to label and add them to the set of training examples. Screen captures of the interface are provided in Supplementary Figure 3.

**Figure 1.**
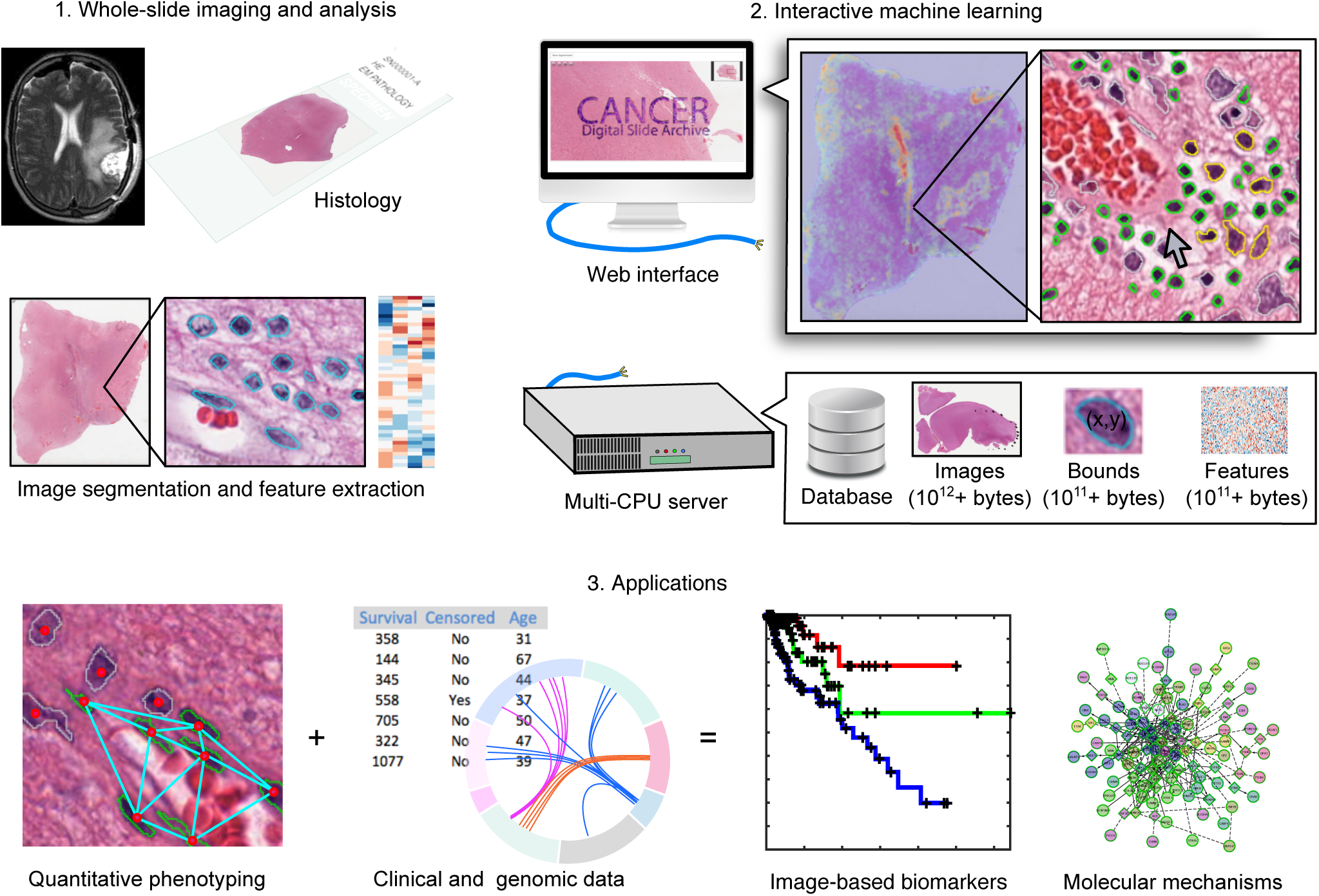
An interactive machine-learning framework for phenotyping histology images. Digitized whole-slide images of tissue sections can be analyzed to extract features describing the shape, texture and staining characteristics of histologic structures like cell nuclei. We created a software framework that enables experts to identify important histologic elements like tumor infiltrating lymphocytes or vascular endothelial cells in these images through interactive training of machine learning classifiers. A browser-based interface provides point-and-click interaction with datasets containing 10^8^+ objects for training classification rules. A multi-CPU server manages the images and boundary and feature data and provides the computational power for visualization and analysis. Classifications generated with this framework can be used to describe the phenotypes associated with cancer-related processes like angiogenesis and lymphocytic infiltration, and to investigate phenotype-genotype associations and phenotypic prognostic biomarkers.

The active-learning methods used in classification rule training are illustrated in Figure 2, using classification of tumor-infiltrating lymphocytes as an example. When making a prediction, many classification rules also produce a confidence measure that represents the expected accuracy of this prediction. In active learning, low confidence objects are labeled to fill gaps in the training set to improve accuracy. Given a classification rule, the set of unlabeled objects are first classified to generate prediction confidences. Labels are then solicited for low confidence objects, and these objects are added to the training set to re-train the classification rule. Figure 2A illustrates a classification rule as a partition of the histomic feature space into region corresponding to distinct cytologic classes. Feature values determine the positions of objects in this space, with classifications being less certain approaching the partition boundary. By iterating between labeling low confidence objects and re-training the classification rule, a feedback loop is established with the user to build a comprehensive training set that increases expected prediction accuracy. This label-update-predict cycle is repeated until the desired performance is achieved (see Supplementary Figure 4 and online Methods for random forest implementation).

**Figure 2.**
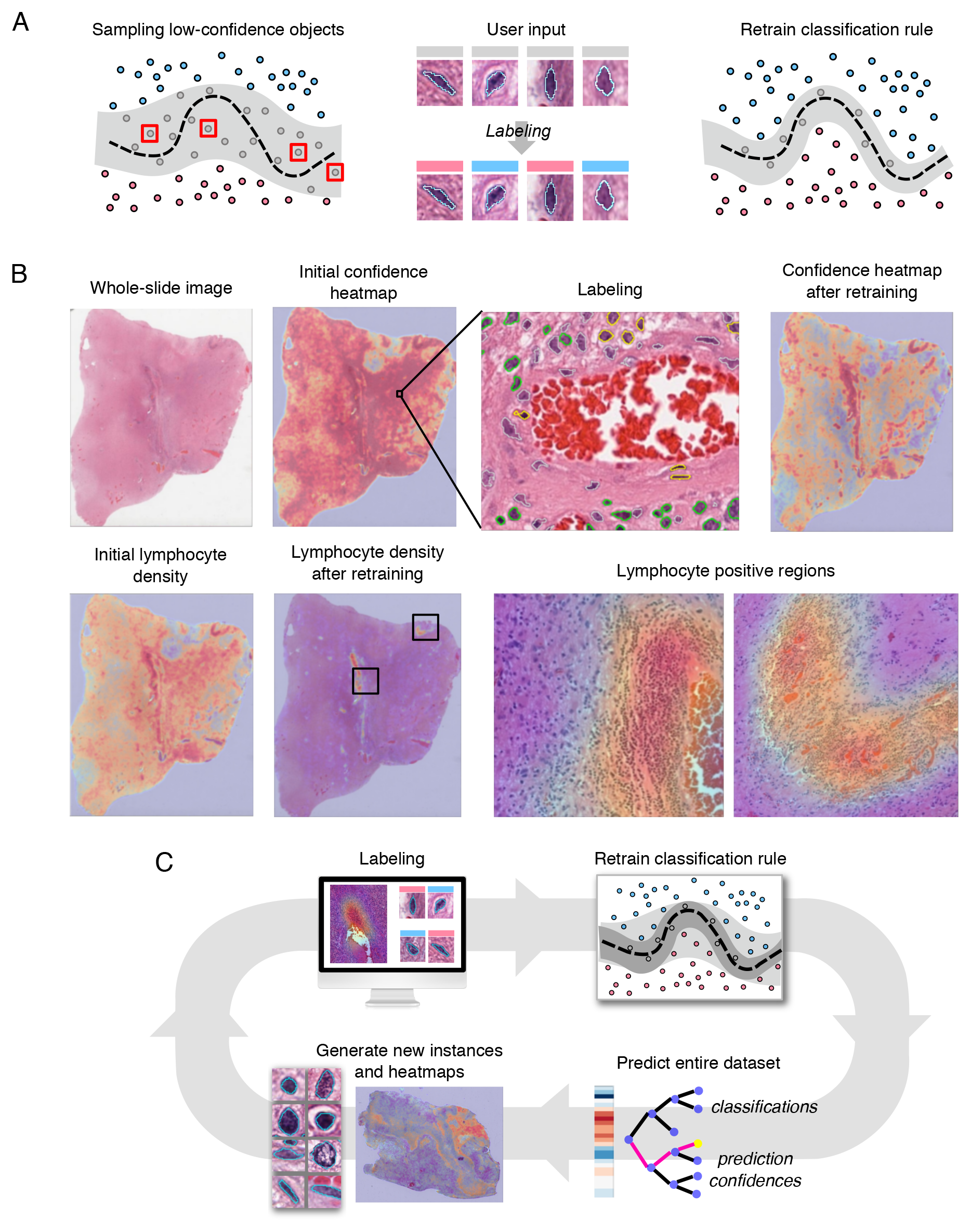
Active learning for efficient classification rule training. (A) (Left) A classification rule aims to learn an unknown decision boundary (black) that separates classes of objects in feature space. A margin (gray) surrounding this boundary contains objects with low prediction confidence that are difficult for the rule to classify. (Center) Instance-based learning presents unlabeled low-confidence objects to the user for labeling. (Right) Retraining the classification rule with these labels shrinks the margin towards the decision boundary improving classification accuracy. (B) Heatmap-based learning directs users to image regions that are enriched with lowconfidence objects for labeling. (Top) Correcting prediction errors (yellow) in low-confidence regions (red) and retraining reduces the number of low-confidence objects. (Bottom) Classification rule specificity is improved by re-training. Here the heatmaps indicate the density of cells positively classified as lymphocytes before and after retraining. (C) Active learning is an iterative process: the user first labels objects guided by active learning, then the classification rule is retrained and applied to the entire dataset, and lastly new instances and heatmaps are generated.

HistomicsML uses two active learning methods to solicit labels: 1. Instance-based and 2. Heatmap-based. Instance-based learning presents the user with 8 of the least confident objects with and array of thumbnail images that can be labeled. Clicking an instance/thumbnail will direct the whole-slide viewer to the location of this object so that the surrounding tissue context can also be visualized. In heatmap-based learning, heatmaps representing prediction confidence are generated for each whole-slide image, enabling users to identify and zoom into “hotspot” regions that are enriched with low-confidence objects for rapid labeling. Slides with hotspots can be identified using an image gallery where slides are sorted based on aggregate confidence statistics.

### Rapid and accurate classification of vascular endothelial cells in gliomas

We used HistomicsTK (http://github.com/DigitalSlideArchive/HistomicsTK) to generate features for 360 million cell nuclei using 781 images (464 tumors) from The Cancer Genome Atlas Lower Grade Glioma (LGG) project. We trained a classification rule to identify vascular endothelial cell nuclei (VECN) and validated its performance using 67 slides not used in training (see Figure 3, Supplementary Figure 5). The VECN classifier was initialized by manually labeling 8 nuclei, and refined with both instance-based and heatmap-based learning to label 135 nuclei in 27 iterations. The VECN classifier is highly sensitive and specific, achieving an area-under-curve (AUC) of 0.964 and improving over the initial rule with AUC=0.9234.

**Figure 3.**
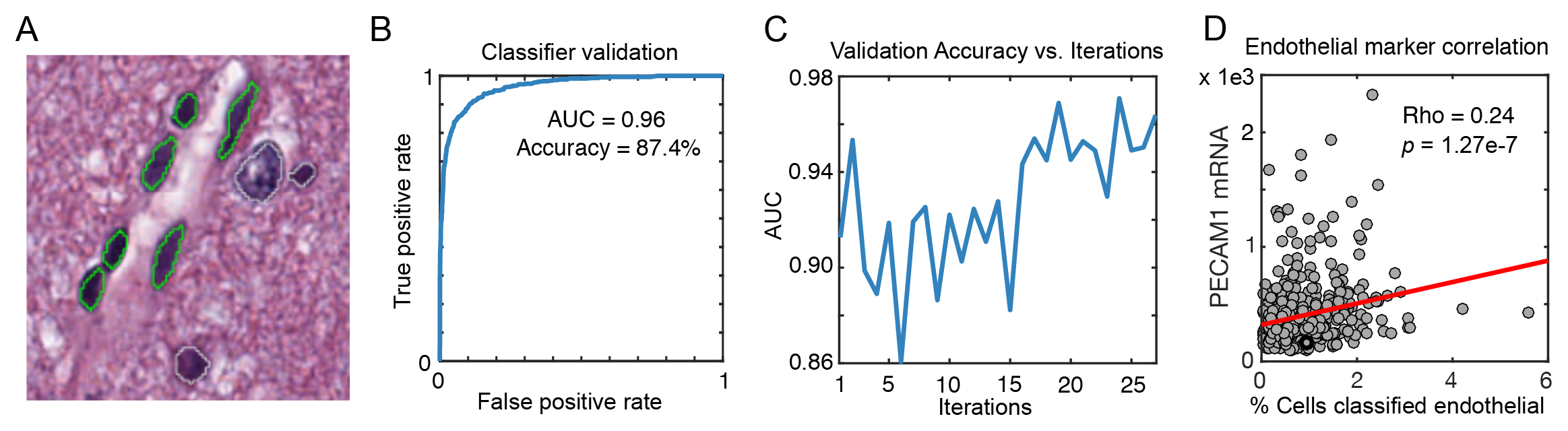
Classifying vascular endothelial cells in gliomas. **(A)** We used active learning to train a classification rule to identify vascular endothelial cell nuclei in lower-grade gliomas (highlighted in green). **(B)** Prediction rule accuracy was evaluated using area-under-curve (AUC) analysis. **(C)** AUC was evaluated at each active learning iteration to measure improvement in prediction accuracy. **(D)** For additional validation, we correlated the percentage of positively classified endothelial cells in each sample with mRNA expression levels of the endothelial marker *PECAM1* using measurements from TCGA frozen specimens (image analysis measurements were performed in images of formalin-fixed paraffin embedded sections from the same specimens).

To further validate our VECN classifier, we correlated mRNA expression of the endothelial marker *PECAM1* with the fraction of cells classified as VECs in each specimen. *PECAM1* expression was significantly positively correlated with Percent-VECN (Spearman rho=0.24, **p=1.27e-7**). We note that the mRNA measurements originate from frozen materials where image analysis was performed on fixed and paraffin embedded tissues that originate from same primary tumor but with unknown proximity to the mRNA sample.

To evaluate system responsiveness, we measured the time required for the update-predict cycles. We evaluated various sized datasets ranging from 10^6^-10^7^ objects (see Supplementary Table 2). We observed a consistent linear increase of 1 second per 5.5 million objects on our 24-core server. This translates to a 10 second learning cycle for a 50 million-object dataset.

### Phenotyping microvascular structures in gliomas

After demonstrating accurate VECN classification, we developed and validated quantitative metrics to describe the phenotypes of microvascular structures (see Figure 4). Gliomas are among the most vascular solid tumors, with microvascular structures undergoing apparent transformations in response to signaling from neoplastic cells. *Microvascular hypertrophy*, or thickening of microvascular structures, represents an activated state where endothelial cells exhibit nuclear and cytoplasmic enlargement due to increased transcriptional activity. *Microvascular hyperplasia* represents the accumulation, clustering and layering of endothelial cells due to their local proliferation. While microvascular changes are understood to accompany disease progression, the prognostic value of quantitating their phenotypes in gliomas has not been established in the era of precision medicine, and may be beyond the capacity of human visual recognition.

**Figure 4.**
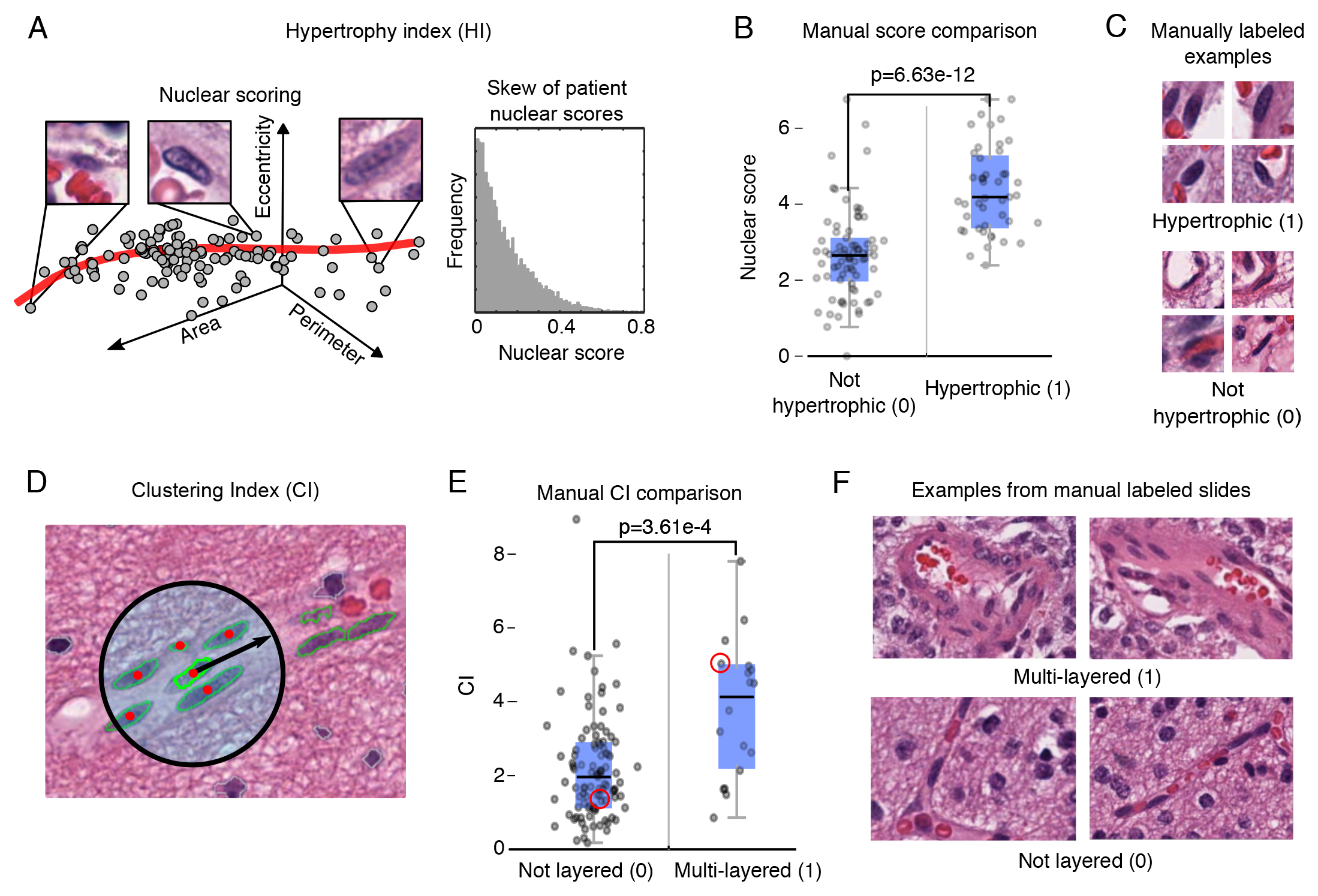
Quantitative phenotyping of microvasculature in gliomas. Microvascular structures undergo visually apparent changes in response to signaling within the tumor microenvironment. **(A)** We measured nuclear hypertrophy using a nonlinear curve to model the continuum of VECN morphologies. A hypertrophy index (HI) was calculated for each patient to measure the extremity of VECN nuclear hypertrophy score values. **(B)** We validated nuclear scores using nuclei that were manually labeled nuclei as hypertrophic / non-hypertrophic. **(C)** Examples of cell nuclei used in validation. **(D)** We implemented a clustering index (CI) to measure the spatial clustering of VECN as a readout of hyperplasia. CI measures the average number of VECN within a 50-micron radius of each VECN in a sample. **(E)** CI was compared to manual assessments of hyperplasia a multi-layered / not layered. **(F)** Example microvascular structures from two of the slides used in validating CI.

Nuclear hypertrophy was scored using a nonlinear model to represent the continuum of VECN morphologies (see Methods). Nuclear scores were validated using 120 manually labeled VECN (45 hypertrophic, 75 non-hypertrophic) to show that nuclei labeled as hypertrophic score significantly higher (Wilcoxon **p=8.63e-12**). A hypertrophy index (HI) was then calculated to summarize hypertrophy at the patient level (see Methods). Hyperplasia was measured using a clustering index (CI) to capture the extent of proliferation and spatial clustering of VECN. CI was calculated at the patient level as the average number of VECN within a 50-micron radius centered at each VEC nucleus. CI was also compared to manual slide-level assessments of microvascular proliferation in 137 slides (18 presenting a multilayered phenotype) to show that images where multilayered structures are present associate with higher CI values (Wilcoxon **p=3.61e-4**).

### Microvascular phenotypes predict survival

Diffuse gliomas are the most common adult primary brain tumor and are uniformly fatal. Survival of patients diagnosed with infiltrating glioma depends on age, grade and molecular subtypes that are defined by *IDH* mutations and co-deletion of chromosomes 1p and 19q (22). The lower grade gliomas (grades II, III) exhibit remarkably variable survival ranging from 6 months to 10+ years. Aggressive IDH wild-type (IDHwt) gliomas having an expected survival of 18 months, where patients with gliomas having IDH mutations and 1 p/19q co-deletions (IDHmut-codel) can survive 10+ years. Gliomas with IDH mutations but lacking codeletions (IDHmut-non-codel) have intermediate outcomes with survival ranging from 3-8 years. The accuracy of grade in predicting outcomes varies depending on subtype (23).

We first investigated associations between hyperplasia and hypertrophy, grade and molecular subtype in the TCGA cohort using CI and HI (see Figure 5A and Supplementary Table 1). We found that IDHwt gliomas exhibit a greater degree of microvascular hyperplasia than the less aggressive subtypes (Kruskal-Wallis **p=8.43e-6**), and that increased hyperplasia is also associated with higher grade within each molecular subtype (Wilcoxon IDHwt **p=4.99e-4**, IDHmut-non-codel **p=1.96e-6**, IDHmut-codel **p=2.08e-4**). While differences in microvascular hypertrophy across subtypes and grades were not statistically significant (Wilcoxon **p=0.747**), the median HI for grade III disease was higher within each subtype. We also explored subtypes by using median CI or HI values to stratify patients into high/low risk groups (see Supplementary Figure 6). Kaplan-Meier analysis found these “digital grades” were marginally prognostic in IDHmut-codel gliomas (log-rank CI **p=6.87e-2**, HI **p=5.09e-2**) and IDHwt gliomas (CI **p=4.68e-2**), but remarkably neither CI nor HI could discriminate survival in the IDHmut-non-codel gliomas. Similar discrimination patterns were observed when stratifying by WHO grade.

**Figure 5.**
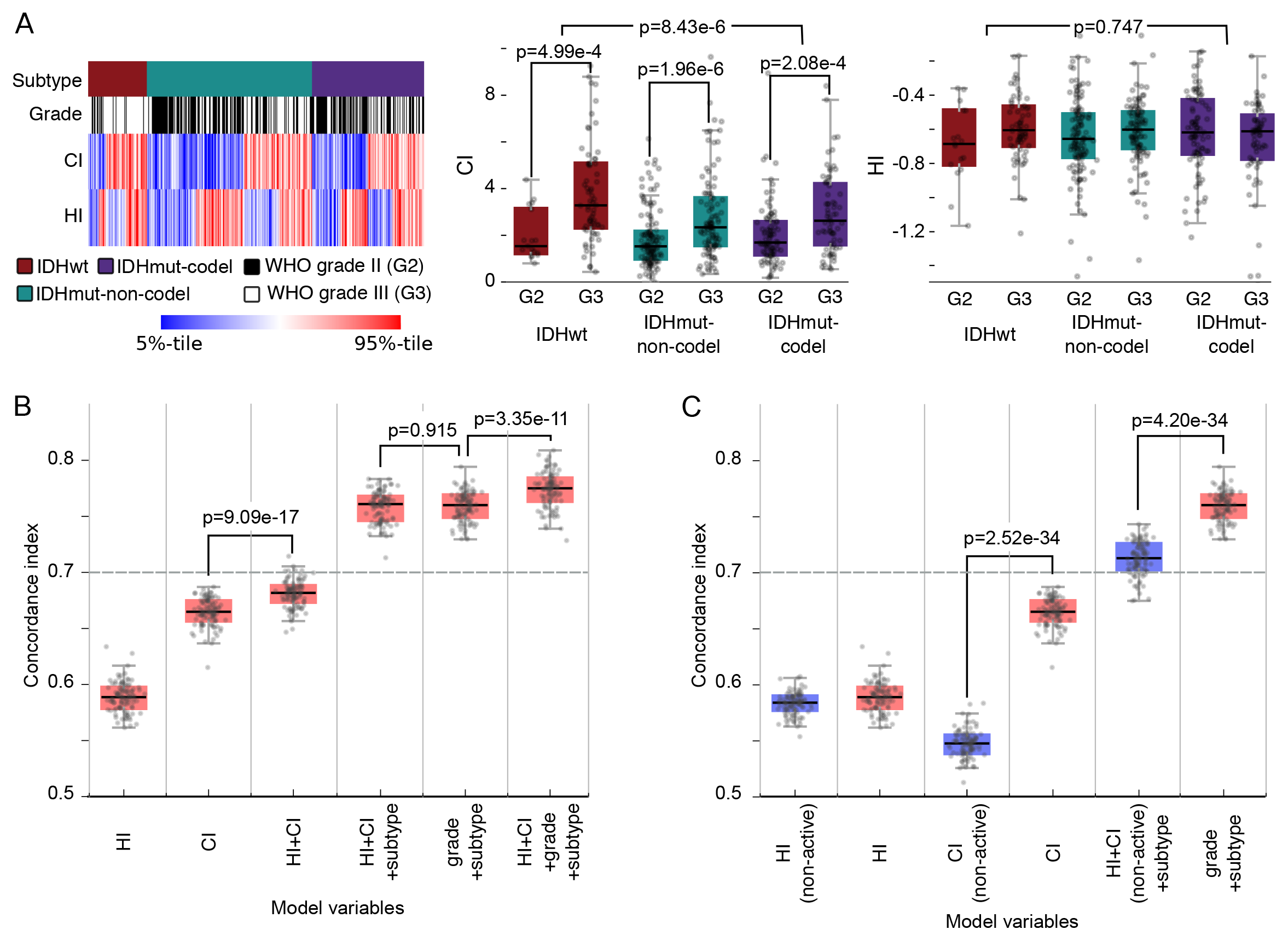
Predicting survival of glioma patients with microvasculature phenotypes. **(A)** HI and CI were compared with important clinical metrics including WHO Grade and molecular subtype. **(B)** We trained cox hazard models using combinations of phenotypic and clinical predictors to assess their prognostic relevance and independence. Models were trained and evaluated using 100 randomizations of samples to training/testing sets. The dashed line represents the c-index corresponding to molecular subtype in this cohort. **(C)** We compared the accuracy of models based on HI and CI generated using a classifier trained with active learning (red) with HI and CI generated using a standard classifier trained without active learning (purple).

After investigating associations with grade and subtype, we used a modeling approach to evaluate the prognostic value of microvascular phenotypes. Cox hazard models were created with various combinations of predictors including grade, subtype, CI and HI (see Figure 5B). Patients were randomly assigned to 100 nonoverlapping training / validation sets, and each was used to train and evaluate a model using Harrell’s concordance index (see Methods). Although HI-only models perform only slightly better than random (median c-index 0.58), HI+CI models perform significantly better than CI-only models (**p=9.09e-17**). HI+CI provide prognostic value independent of molecular subtype, improving the subtype c-index from 0.70 to 0.76. HI+CI also performs as well as grade when combined with subtype (**p=0.915**), even though grade incorporates many more histologic criteria than microvascular appearance. Finally, HI+CI also have prognostic value independent of grade+subtype, increasing median c-index to 0.78 (Wilcoxon **p=3.35e-11**).

### Active learning training improves prognostication

To evaluate the benefit of active learning training, we repeated our experiments using a classification rule trained with a standard approach where the expert constructs a training set without the aid of active learning feedback. Using the same image collections described above, 135 cell nuclei were labeled in the training images (roughly evenly split between VECN and non-VECN). A classification rule was trained using these labels and applied to the dataset to compare classification and prognostic modeling accuracies with the active learning classifier.

The validation AUC of the standard classifier was 0.984 (AUC=0.964 for active learning classifier). While the AUC measured on the validation set was higher, the standard learning classifier is much less specific on the entire dataset, producing very high estimates of percent-VECN in the TCGA cohort ranging from 7.1-57.2% (compared to 0.02–5.6% percent-VECN for active learning). Agreement between *PECAM1* expression and percent-VECN was much lower for the standard classifier percent-VECN (Spearman rho=0.16 versus 0. 24). We calculated updated HI and CI metrics using the standard classifier results and found that prognostic models based on these metrics were no longer predictive of survival (see Figure 5C). The median c-index of models based on CI alone fell to <0.55 (Wilcoxon **p=2.52e-34**). Models incorporating HI+CI+subtype were also no longer equivalent to subtype+grade models (**p=4.20e-34**), and only slightly better than subtype.

### Integrating phenotypic measures with genomic information

The molecular mechanisms of angiogenesis in gliomas have been studied extensively, and are targeted through anti-VEGF therapies like Bevacizumab (24). To investigate the molecular pathways associated with CI/HI, we performed gene-set enrichment analyses (25) to correlate CI and HI with mRNA expression. We analyzed IDHwt and IDHmut-codel gliomas separately since mechanisms may vary across subtype (IDHmut-non-codel gliomas were not analyzed). A partial list of pathways enriched at FDR q<0.25 significance is summarized in Table 1 (extended results in Supplementary Table 3).

**Table 1.** Molecular pathways enriched with phenotype-correlated transcripts. Gene set enrichment analysis of the correlations between HI / CI and gene expression identified multiple pathways associated with gliomas and vascularization. Many of the significantly enriched pathways are specific to one molecular glioma subtype. Extended results are presented in Supplementary Table 2.

| Pathway Group | Pathway name | Leading-edge genes | Subtype / metric (directionality) | Nominal p-value (FDR q-value) |
| --- | --- | --- | --- | --- |
| Classical angiogenesis pathways | * HIF1-alpha transcription factor | *PFKL, PFKFB3, ALDOA, PGK1, HK1* | IDHmut-codel / HI (+)<br>IDHmut-codel / CI (-) | 0.033 (0.179)<br>< 0.001 (0.116) |
|  | HIF2-alpha transcription factor | *VEGFA, VHL, ARNT* | IDHwt / CI (+)<br>IDHmut-codel / CI (+) | 0.004 (0.017)<br>0.024 (0.116) |
|  | VEGFR1/2 mediated signaling | *BRAF, MAPK1/14* | IDHwt / HI (+)<br>IDHmut-codel / CI (+) | 0.012 (0.144)<br>0.009 (0.12) |
|  | * VEGFR1 specific signals | *MAPK1, NRP1/2* | IDHwt / HI (+) | 0.007 (0.19) |
|  | Angiopoietin receptor TIE-2 mediated signaling | *MAPK1/14, NFKB1, PIK3C* | IDHwt / HI (+)<br>IDHwt / CI (+)<br>IDHmut-codel / CI (+) | 0.014 (0.147)<br>0.015 (0.063)<br>0.009 (0.087) |
|  | * PDGFRA signaling | *PIK3CA, FOS, PDGFRA* | IDHmut-codel / HI (+) | 0.014 (0.08) |
| Developmental signaling pathways | Notch signaling network | *NOTCH1, MAML1/2, MYC* | IDHwt / CI (+)<br>IDHmut-codel / CI (+) | < 0.001 (< 0.001)<br>0.02 (0.146) |
|  | * Notch mediated HES/HEY network | *HEY1, NOTCH1, MAML1/2, HIF1A* | IDHwt / HI (+)<br>IDHwt / CI (+) | 0.007 (0.143)<br>< 0.001 (< 0.001) |
|  | * WNT signaling | *WNT3A, GSK3B* | IDHmut-codel / CI (+) | 0.021 (0.085) |
|  | * Regulation of nuclear beta catenin signaling |  | IDHmut-codel / CI (+) | 0.004 (0.054) |
|  | * GLI-mediated Hedgehog signaling |  | IDHwt / CI (+) | 0.015 (0.063) |
| Other pathways | * Regulation of SMAD2/SMAD3 signaling | *SMAD3/4, MAPK1, MAP3K1* | IDHwt / HI (+)<br>IDHwt / CI (+) | 0.002 (0.008)<br>0.024 (0.139) |
|  | * SMAD2/SMAD3 nuclear signaling | *SMAD3/4, CDK2/4, CDKN1A, AKT1, MYC* | IDHwt / CI (+) | < 0.001 (< 0.001) |
|  | * FOXM1 transcription factor network | *FOXM1, GSK3A, MYC, FOS* | IDHmut-codel / CI (+) | < 0.001 (< 0.001) |

Given the association between angiogenesis and hypoxia, we anticipated pathway analysis to identify strong relationships between microvascular phenotypes and classic hypoxia and metabolic glycolysis pathways. We found both HIF2A and VEGFR1/2 mediated signaling pathways were both upregulated with increasing CI and HI. Among the most strongly correlated genes were those involved in hypoxia and angiogenesis including *VEGFA*, *VHL*, *ARNT*, *PGK1* (26), *ADM* (27), and *EPO*, as well as glycolytic response mediators *HK1*, *PGK1*, *ALDOA*, *PFKFB3*, *PFKL* and *ENO1*. Angiopoietin receptor (28) and Notch signaling (29) pathways were also significantly enriched in both glioma subtypes.

Pathways with enrichment specific to IDHwt gliomas included Notch mediated regulation of *HES/HEY* (30), GLI-mediated hedgehog signaling (31), and SMAD signaling (32), all of which have been linked to angiogenesis or regulation of structure and fate in vascular endothelial cells. Pathways with enrichment specific to IDHmut-codel gliomas included WNT and beta-catenin signaling, and PDGFRA signaling (PDGFRA amplification is frequent in IDHmut-codel gliomas).

We note that angiogenesis generally accompanies disease progression in gliomas, and that pathway enrichments may reflect molecular patterns associated more generally with disease progression in addition to angiogenesis-related microenvironmental signaling.

## DISCUSSION

HistomicsML addresses the unique challenges presented by the scale and nature of whole-slide imaging datasets to enable investigators to extract phenotypic information. It is open-source and is available as a software container for easy deployment.

The endothelial cell classifier trained with active learning was highly accurate, despite labeling only 135 cell nuclei (AUC=0.9643). The visualization and learning capabilities of HistomicsML simplifies the training by enabling experts to rapidly label objects and to re-train and review classification rules in seconds. The web-based interface provides remote access to terabyte datasets, and enables fluid and seamless display of image analysis boundaries and class predictions associated with 10^8^+ histologic objects. Active learning directs labeling efforts by guiding users to examples that provide the most benefit for classifier training, and improves efficiency by avoiding labeling of redundant examples.

Phenotypic metrics obtained using our endothelial classifier were validated using human annotations, and able to accurately predict survival of lower-grade glioma patients. We identified significant associations between microvascular phenotypes, grade, and recently defined molecular subtypes of gliomas. These investigations are timely in the current era of precision medicine, in which prognostic biomarkers have not been established within newly emergent genomic classifications of cancers. While it has long been established that angiogenesis is related to disease progression in gliomas, we showed that HistomicsML can be used to precisely measure subtle changes in microvasculature that perform as well as grade in predicting survival. Active learning training was shown in these experiments to both improve prognostication and agreement between histologic and molecular markers of VECNs. Integrating phenotypic metrics with genomic data identified recognized molecular pathways associated with angiogenesis and disease progression. These analyses are a template for how HistomicsML can link histology, clinical and genomic data to explore the prognostic and molecular associations of histologic phenotypes in other diseases. Histology contains important information that can be difficult or impossible to ascertain through genomic assays. Recent developments in the deconvolution of gene expression data can accurately estimate the proportions of cell types in a sample, but these approaches cannot provide spatial or morphologic information that often contains considerable prognostic or scientific value.

Future development of HistomicsML will focus on enabling the classification of image-regions and improved scalability for larger datasets. Classification of image regions characterized by texture analysis or autoencoders can address the classification of complex multicellular structures while avoiding the need for explicit segmentation and is a simple extension of HistomicsML. The memory footprint of feature data is currently a limitation that prevents HistomicsML from scaling to larger datasets. We plan to improve memory management, and to utilize commodity graphics processors to enable better scalability.

## METHODS

### Data

Whole slide images, clinical and genomic data were obtained from The Cancer Genome Atlas via the Genomic Data Commons (https://gdc.cancer.gov/). Images of formalin-fixed paraffin-embedded “diagnostic” sections from the Brain Lower Grade Glioma (LGG) cohort were reviewed to remove images of sections with tissue processing artifacts including bubbles, section folds, pen markings and poor stain quality. For this paper a total of 781 whole-slide images were analyzed. Genomic data (described below) was derived from frozen materials from the same specimens. The relationship of diagnostic sections and frozen materials is unknown, other than that they originate from tissues produced during the same surgical resection.

### Image analysis segmentation and feature extraction

The software pipeline used to segment cell nuclei and measure their histomic features is shown in Supplementary Figure 1. This pipeline utilizes algorithms provided by the HistomicsTK Python library for histologic image analysis (http://github.com/DigitalSlideArchive/HistomicsTK) to perform color normalization, nuclear masking and splitting, feature extraction, and database ingestion. Each whole slide image was normalized to a standard H&E with desirable color characteristics using Reinhard normalization. Tissue pixels were first masked from the background using linear discriminant analysis and then the mean and standard deviation of the tissue pixels in the L*A*B color space were calculated. These moments were mapped to match the moments of a color standard image prior to inversion back to RGB color space. This color normalization process considerably improves the quality of subsequent image analysis steps, improving the consistency of segmentation results and image features. Whole slide images were tiled into 4096 x 4096 pixel tiles and processed independently to highlight cell nuclei a color deconvolution algorithm was first applied to digitally separate the hematoxylin and eosin stains. The deconvolved hematoxylin intensity images were then masked to identify pixels corresponding to cell nuclei using a combination of adaptive thresholding and morphological reconstruction to remove background debris. Closely packed nuclei were then split using a watershed segmentation applied to the laplacian-of-gaussian response of the hematoxylin image. Each nucleus was described using a set of 48 features describing its shape, intensity and texture. These features include eccentricity, solidity and fourier shape descriptors (shape), statistics of hematoxylin signal including variance, median, mean, min/max, kurtosis, skew and entropy (intensity) and statistics of hematoxylin intensity gradients (texture). Color normalization, segmentation and feature extraction are carried out in a cluster-computing environment using Torque to distribute slides to different computing nodes. Boundaries generated by segmentation are stored in a text-delimited format and ingested into a SQL database to drive visualization in the web-based interface. Features are stored as matrices in a structured HDF5 format and stored on a RAID array along. Prior to machine learning analysis all features are z-scored using the aggregate mean and standard deviation calculated across the feature profiles of every cell in the dataset.

### Validation and training

We selected 67 digital slides from the LGG cohort to serve as a validation set for measuring the performance of a vascular endothelial classifier. In each slide, we selected a field containing a mixture of nuclei from vascular endothelial cells and other cell types (tumor nuclei and inflammatory cells, for example). Each correctly segmented nucleus in the field was labeled as either vascular endothelial or other. Incorrectly segmented nuclei and nuclei that were too ambiguous to classify with a high degree of certainty were ignored. In total 2479 cell nuclei were labeled. Each annotation was reviewed by a board-certified neuropathologist using our review function. Classifiers used for reporting accuracy results were trained using a mixture of instance-based and heatmap-based review.

Labels to validate hypertrophy index and clustering index were acquired by manual inspection of digital slide images by a board-certified pathologist who was blinded to the computer-generated HI and CI scores. A selection of 120 cell nuclei classified as VECN were labeled as either hypertrophic (45 nuclei) or non-hypertrophic (75 nuclei) using our the HistomicsML Review tool. Nuclear hypertrophy scores were compared for these manually labeled nuclei using a non-parametric Wilcoxon sign rank test. For clustering index, 137 slides were manually reviewed to determine if they present microvascular hyperplasia and proliferation (multilayered vessels). CI scores were compared for slides containing multi-layered vessels and slides not containing multilayered vessels using the Wilcoxon test.

### Software

All source code for the active learning system are published under a Apache 2.0 license at (https://github.com/cooperlab/ActiveLearning). In addition to source, we have published a software container at (https://hub.docker.com/r/histomicsml/active/) for easy deployment. This Docker container comes with sample data loaded from a single whole-slide image.

### Machine learning

The active learning paradigm is agnostic to the specific choice of classifier, and any classifier that provides a measure of prediction confidence can be utilized. In our implementation we chose random forests due to their resistance to overfitting, computational efficiency and simplicity in design (see Supplementary Figure 4). We used the implementation provided by the OpenCV (v2.4.10) machine learning library. The parameters for the random forest implementation are the number of trees (fixed at 100), each tree with a fixed maximum depth of 10, and a variable number of features selected for node splits (calculated as the floor of the square root of number of histomic features for each object). The prediction confidence for each object is calculated by voting on the predictions of the trees. Prediction confidence for a random forest classifier evaluated on object *i* is calculated as

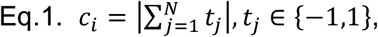

where *t_j_* is the prediction tree *j* of *N* total trees. Maximum prediction confidence is achieved if all trees are in consensus on the predicted class. Minimum confidence is achieved if the trees are evenly split. When applying prediction to tens or hundreds of millions of objects, multiple threads are spawned on the server to minimize user delay and to maintain system responsiveness.

### Endothelial clustering index

CI was calculated using a spatial statistic based on a modified version of Ripley’s K-function (36). Ripley’s K-function, originally developed for geographic and epidemiologic applications, captures the “degree of spread” of events in spatial domains. Since microvascular hyperplasia and proliferation are, by definition, associated with increases in VECN density, we excluded Ripley’s density normalization in our proliferation metric. We also ignored edge-effect corrections due to the extremely large number of objects and the relative rare scarcity of objects at the edge of tissue sections. CI was calculated as:

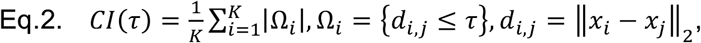

where *K* is the number of objects/nuclei in the image, *d_i,j_* is the Euclidean distance between objects *i*, *j*, *τ* is a positive search radius. This effectively calculates the average number of objects within distance *τ* of each object in the slide. In our experiments, CI was calculated for objects positively classified as VECN with a search radius of *τ* = 50-microns. KD tree indexing was used to accelerate the computational search for neighboring VECN.

### Endothelial hypertrophy index

The first step in calculating HI is to score the morphology of individual nuclei classified as VECN. A principal curve was trained to model the continuum of VECN morphologies and then used to score the extent of hypertrophy of each VECN nucleus (37). The principal curve was used to model the histomic features of VECN with a one-dimensional nonlinear curve, modeling the morphologic continuum of from small, thin normal appearing vascular endothelial nuclei to enlarged hypertrophic nuclei that are reactive to the surrounding tumor microenvironment. The principal curve models the feature vector values *f_i_* of object *i* as

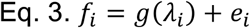

where *g* is a 1D nonlinear curve in feature space parameterized by the free variable I, and e is a random component. After fitting the principal curve, each nucleus is scored by projecting its feature profile onto the curve, and calculating the path length from the projection to the curve origin

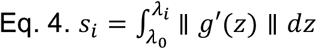

where *λ_0_* is the origin of the curve, *λ_i_* is the location of the least-squares projection of *f_i_* onto the curve, and *g*’ is the curve tangent function. Nuclei that are more hypertrophic in the morphologic continuum will have longer path length values and thus higher nuclear scores. Our analysis constructed the principal curve using histomic shape features for nucleus area, eccentricity and perimeter. The directionality for the beginning/end of the curve was established by initializing the curve fitting with a single normal appearing nucleus and a single hypertrophic nucleus.

At the patient-level, HI was calculated to represent the population-level skew of VECN towards more hypertrophic morphologies. HI was measured as the negative skew of nuclear scores

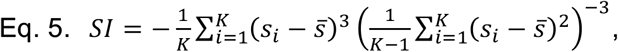

where *K* is the number of objects/nuclei in the image, *s_i_* is the hypertrophy score of object *i*, and 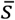 is the mean hypertrophy score. This statistic increases in value as the right tail of the score distribution grows.

### Clinical and genomic data

Genomic and clinical data were acquired using the TCGAIntegrator Python interface (https://github.com/cooperlab/TCGAIntegrator) for assembling integrated genomic and clinical views of TCGA data from the Broad Institute Genomic Data Analysis Center Firehose (https://gdac.broadinstitute.org/). The same genomic platforms were used across all experiments. Gene expression values were taken as RSEM values from the Illumina HiSeq 2000 RNA Sequencing V2 platform. Genomic classifications for IDH/1p19q status were obtained from the Supplementary Material of (38).

### Pathway analysis

Agreement between gene expression and HI / CI was measured using Spearman rank correlation between RNAseq values and CI or HI values. To account for the fact that each subtype and metric may have different molecular mechanisms, we performed four gene set enrichment analyses: IDHwt/HI, IDHwt/CI, IDHmut-codel/HI, and IDHmut-codel/CI. We did not analyze gliomas from the IDHmut-non-codel subtype since microvascular phenotypes were not found to be prognostic within this group. Gene symbols were filtered to remove entries not present or aliased in the HUGO Gene Nomenclature Database (http://www.genenames.org/) (39). Spearman correlations were used to construct an .rnk file for each of the four analysis scenarios described above, and used to perform a Gene Set Enrichment Analysis using the GSEAPreranked (v4.2) module in GenePattern with 1000 permutations. We tested enrichment for pathways described in the NCI/Nature Pathway Interaction Database (PID) using a version of the MSigDB (http://software.broadinstitute.org/gsea/msigdb) C2 Curated Gene Sets that was filtered to remove non-PID pathways. We reported both the nominal p-values as well as FDR-corrected q-values produced by GSEA in Table 1 and Supplementary Table 2.

### Statistical testing and prediction performance measures

Variations in HI/CI across molecular subtypes and grade were evaluated using the Wilcoxon test where the independent variable is binary (i.e. grade) or the Kruskal-Wallis test for multi-category independent variables (i.e. molecular subtype). Survival differences in cohorts were evaluated using the log-rank test. Classifier performance was reported as area under receiver operating characteristic curve, calculated on the signed prediction confidence values *p_i_* generated by the random forest classifier. Classification accuracy was reported using a zero-threshold on prediction confidence values. The performance of prognostic models was reported using Harrell’s concordance index (40).

### Computing hardware

All studies were performed using a multi-socket multicore server equipped with two Intel Xeon e5-2680 v3 2.5GHz processors (total 24 cores / 48 threads), 128 GB memory, and 14 TB main disk storage in a RAID10 array.

### Data availability

This paper was produced using large volumes of publicly available image data. The authors have made every effort to make available links to these resources as well as making publicly available the software methods used to produce these analyses and summary information. All data not published in the tables and supplements of this article are available from the corresponding author on request.

## ACKNOWLEDGEMENTS

This work was supported by U.S. National Institutes of Health, National Library of Medicine Career Development Award K22LM011576, and National Cancer Institute grant U24CA194362, and with funds from the Emory Winship Cancer Institute.

## AUTHOR CONTRIBUTIONS

M.N., S.L., D.A.G. and L.A.D.C. developed the software framework and algorithms and performed documentation and installation packaging. M.A. and L.A.D.C. performed analysis of clinical and genomic data. J.V.V. and D.J.B. provided neuropathology review and inputs on the design of phenotyping metrics, and assisted with the interpretation of results. D.A.G., M.N., M.A., S.L., S.H.H., J.V.V., D.J.B. and L.A.D.C. assisted with manuscript development. L.A.D.C. conceived of the software concept and supervised the project.

